# An Amyloid beta-derived Septapeptide has hMCP-1 Chemokine Activity

**DOI:** 10.1101/424838

**Authors:** Diane Van Alstyne

## Abstract

Septapeptides (“septas”), previously identified as meningitis-specific antigens, defined by a rubella virus (RV) monoclonal antibody, were found in human Monocyte Chemoattractant Protein (hMCP-1) and on the surface of meningitis-causing bacteria, viruses and spirochetes. Some bacterial septas were tested for Ca^+2^ mobilization through receptor-associated heterotrimeric G-protein binding on THP-1 cells, progenitor cells of circulating peripheral macrophages. Certain of the free septas acted on their own as mild agonist of Ca^+2^ mobilization. Their signal transduction activity may be mediated through a single, or a single class, of receptor, but the data do not link this with the MCP-1 receptor on THP-1 cells. These data support the proposals that (1) the septas represent muteins of the MCP-1 active site for stem cell activation to macrophages and (2) infectious organisms may conserve and employ these sequences in order to facilitate their transport through the Blood Brain Barrier (BBB) to infect the Central Nervous System (CNS). Other MCP muteins have since been identified in the Alzheimer’s Disease (AD)-associated agents, amyloid beta (Abeta) and prions, as well as in viruses like HIV, known to establish chronic infections in the CNS. Rat glial progenitor cells in tissue culture were used to test for hMCP-1 activity in septas derived from amyloid beta. Nanomolar concentrations of the amyloid beta septa ^13^HHQKLVF^19^ were found to transform more than 60% of the progenitor cells into microglia in tissue culture. Taken together, these data support the hypothesis that amyloid beta may accumulate to increase the number of microglia available to combat chronic infection in the CNS. A new paradigm for neurodegenerative disease is presented.

## Introduction

Models of bacterial meningitis pathogenesis are summarized in several comprehensive reviews [1-3]. Mechanisms used by microbial pathogens to enter the CNS are divided according to the cellular route involved and whether the organisms breach endothelial cells of the BBB or specialized epithelial cells of blood-choroid barriers. The meningitis causing bacteria (MCB) are commonly referred to as transcellular (passing directly through cells) [1-3], paracellular (passing between cells, through tight junctions)[1-3] and leukocyte-facilitated [4]. The latter was first reported when *S.suis* was observed to use this mechanism to cross the choroid plexus in pigs [5]. The hallmark of this mechanism of infection is exemplified by *L. monocytogenes* which uses its p60, and internalin A, and B, to attract and infect Ly-6C^high^ monocytes [6,7]. Other MCB like *E.coli* K1 [8,9], *M.tuberculosis* [10,11] and *S.enterica* serovar *Typhimurium* [12,13] have also been shown to initiate their specific uptake in macrophages, to replicate intracellularly, and to cross epithelial barriers, suggesting that the trafficking of infected phagocytes may be a common mechanism for a variety of MCB.

Previous work on the transport of meningitis-causing organisms through the BBB identified a series of meningitis-related homologous antigenic sequences (MRHAS), defined as septapeptides homologous to antigenic sites on the structural RV polypeptides, that are recognized by an anti-RV monoclonal antibody (Mab) from the hybridoma RV-1. Members of the MRHAS family were also found to appear in one variant of the chemokine, human Monocyte Chemoattractant Protein-1 (hMCP-1) [14-16]. It was proposed that the incorporation of the hMCP-1 active site peptide conferred an advantage on bacteria, viruses and spirochetes, attracting and infecting a monocyte subset, enabling them to cross the BBB while contained in macrophages which routinely travel through the membrane tight junctions. The MRHAS peptides have since been renamed “septas”.

The meningitis-derived model described above may be viewed as “platform technology”, with a direct application to our understanding of the pathology of AD. AD is an age-related neurodegenerative disorder characterized by the progressive loss of cognitive and executive functions. The main features of AD are the extracellular deposition of Abeta protein into the extracellular amyloid-plaques [17] and the intracellular accumulation of tau protein in the form of neuofibrillary tangles [18]. The relationship between Abeta deposition and the onset of AD pathology is not clear. Membrane damage is thought to play a central role in AD [19] with phospholipid metabolism implicated as the key variable in pathogenesis [20-23].

Abeta is known to induce microglial and astrocyte cell activation which in turn promotes inflammation through the release of proinflammatory mediators. Microglia and astrocytes are innate immune cells in the brain. They constantly scan the tissue, and respond to pathological signals to encapsulate pathogenic foci and remove apoptic cells and debris, much as peripherally-derived macrophages do. Abeta has also been proposed as toxic to neurons through synaptic damage [24]. Loss of synaptic function is associated with cognitive decline in AD. Microglial populations have recently been shown to remove damaged synapses while other subsets restrict neurodegeneration [25,26]. Abeta has also been shown to be an antimicrobial peptide which accumulates in response to bacterial infection [27].

Tau has also been proposed as a mediator of AD. It is one of the microtubule-associated proteins that promote assembly of tubulin into microtubules. As a structural component of the intracellular cytoskeleton, it is expressed mainly in neurons and is involved in the neurofibrillary tangles and threads characteristic of AD [18,28]. Tau pathologies are found in a range of neurodegenerative disorders and progression of the pathologies occurs through prion-like seed-dependent aggregation [28]. One school of thought suggests that intervention in the conversion of normal tau to abnormal forms and in cell-to-cell transmission of tau may be the key to development of disease-modifying therapies for AD and other dementia-related disorders [28,29].

The prion protein has also been proposed as a mediator of some CNS diseases including AD [30-32]. Prions have been defined as host-encoded cellular proteins, associated with the intracellular cytoskeleton, with the ability to undergo structural isomerization to an infectious form, termed PrPSc [33]. The process is thought to be autocatalytic, with PrPSc serving as a template for refolding of PrpC. Protein aggregation and amyloid formation are central pathologic events in a wide range of neurodegenerative illnesses, including Gerstmann-Straussler-Scheinker (GSS) Disease [34,35], multiple system atrophy (MSA) [30], AD, Parkinson’s Disease and prion diseases such as Scrapie. One specific prion protein mutation, lacking a plasma membrane anchor, has been shown to cause GSS disease, characterized by wide-spread amyloid deposition in the brain and the presence of a short protease-resistent PrP fragment similar to similar peptides found in GSS patients. This PrP 23-88 peptide sequence was found to greatly increase disease progression when administered to mice [36]. These finding strongly suggest that prion-mediated spread of neuron dysfunction in the CNS represents a general biological phenomenon of amyloidosis leading to neurodegeneration.

Rat glial progenitor cells in tissue culture (RG cells) have been employed in this study to detect chemokine activity associated with septas. These indicator cells were established employing intracerebral injections of kainic acid to cause a profound loss of neurons, at the injection site in adult rat brain, used initially as a model for Huntington’s Disease [38,39]. Glial progenitor cells from kainic acid-lesioned brains of adult rats were established and maintained in continuous cell culture. Cells expressed the glial fibrillary acidic protein (GFAP), but were negative for three neuronal markers: choline acetyltransferase, tyrosine hydroxylase and glutamic acid decarboxylase [39]. Treatment of RG cells with 1mM dibutyryl cyclic AMP (dBcAMP) [39] resulted in their transformation into mature glia expressing GFAP-positive cells in 80% of the population and the GC-positive (oligodendrocyte-specific galactocerebroside) cells in less than 10% of the population in passage number 35 [41]. Approximately 15% of the mature cells remained unidentified.

This study brings our earlier work on rubella virus and meningitis to bear on the development of a new approach in our understanding of neurodegeneration.

## Materials and Methods

### Sequence homology analyses

Determination of sequence homologies is detailed in two early patents from 1996, using MacDNAsis, Hitachi Software Engineering [14,15] which employed the Lipman-Pearson FASTA algorithm.

### Measurement of MCP-1 activity in septas using Ca^+2^ mobilization

Chemokine activity was measured quantitatively using the standard Ca^+2^ mobilization.assay described in the Cytokine Handbook [42]. THP-1 cells (ATTC), peripheral macrophage progenitor cells, were loaded with Ca^+2^ indicator dye Indo-1AM and their cytoplasmic Ca^+2^ content was measured during laser excitation. Activity was expressed as Relative Fluorescence Intensity (RFI) over time (seconds).

### Maintenance of RG cells in tissue culture

Chemokine activity was measured qualitatively using maturation of rat glial progenitor cells in continuous culture. Kainic acid monohydrate (1ul) (Sigma), containing 5 nmole kainic acid in 0.9% saline was injected into rat striatum under anaesthia. A single rat was anaesthetized and decapitated 5 days following injection and the brain was harvested immediately and placed in a sterile glass culture dish. The striatum from the injection site was dissected, placed in MEM (Gibco) containing 0.2% glucose, 10% fetal calf serum, 5 units/ml penicillin, 5 ug/ml streptomycin and then further minced using a sterile scalpel. The culture dish was then incubated at 37^0^C in a humidified atmosphere containing 5% CO_2_. Each day for the first 4 days the unattached tissue fragments were removed, washed once in fresh medium and returned to the culture dish until a complete monolayer was formed. The resulting monolayers were then routinely subcultured every 5-6 days after trypsinization (0.25% trypsin for 5 minutes at 22^0^C.) The cells were collected by low speed centrifugation, washed twice with complete growth medium and replated in glass culture dishes. Once the cell line was established in glass dishes, the cells were then adapted later to plastic surfaces.

### diButyrylcAMP (diBcAMP)-induced maturation of RG cells

RG monolayer formed at 3 days following subculture, were treated with 100 uM diBcAMP dissolved in complete medium. Medium was first removed from the monolayers and replaced by the diBcAMP-containing medium. Maturation was observed to begin within 1 hour and cells reverted back to the original stem cell monolayer appearance within 24 hours.

### Identification of microglia and astrocytes in treated RG monolayers

At maximal maturation, RG cells were stained using ThermoFisher Scientific CD11b, GFAP (Molecular Probes) monoclonal antibodies and DAPI (4’,6-diamidino-2-phenylindole dihydrochloride), nucleic acid stain. The Alexa Fluor dye-labeled anti-CD11b (monoclonal antibody M1/70, Alexa Fluor 488 and anti-GFAP (monoclonal antibody 131-17719), antibodies were supplied as a 1mg/ml solution in phosphate buffered saline (PBS), pH 7.2, containing 5mM sodium azide and 0.1% bovine serum albumen (BSA). Antibodies were used at a concentration of 5ug/ml.

### Custom peptide synthesis

Peptides derived from Abeta were provided by Kinexus Techologies, Inc., Vancouver, BC, Canada. Each peptide was provided as 5 mg, freeze-dried, linear peptide with a C-terminus amide and N-terminus free amino acid, at ≥95% purity. Samples were shipped and stored at 4^0^C. Prior to use, they were dissolved in DMSO and then diluted further in PBS.

### Abeta-derived peptide induced maturation of RG cells

Peptides were dissolved in dimethyl sulfoxide (DMSO) and serially diluted in PBS to 10^−7^M. RG monolayers were grown to confluency in 5 ml plastic dishes by day 3 after subculture. On day 3 the medium was removed from each dish and replaced with 1 ml of complete medium containing a single concentration of an Abeta-derived peptide. This was done to maximize the chance for peptide binding to cells. After 1 hour, an additional 4 ml of medium was added. Monolayers were incubated for 72 hours. Each peptide concentration was tested in triplicate.

## Results

The following figure lists several of the MCP-1 -derived septas in some meningitis-causing bacteria, viruses and spirochetes as well as in Abeta and prion proteins.

**Figure 1.**
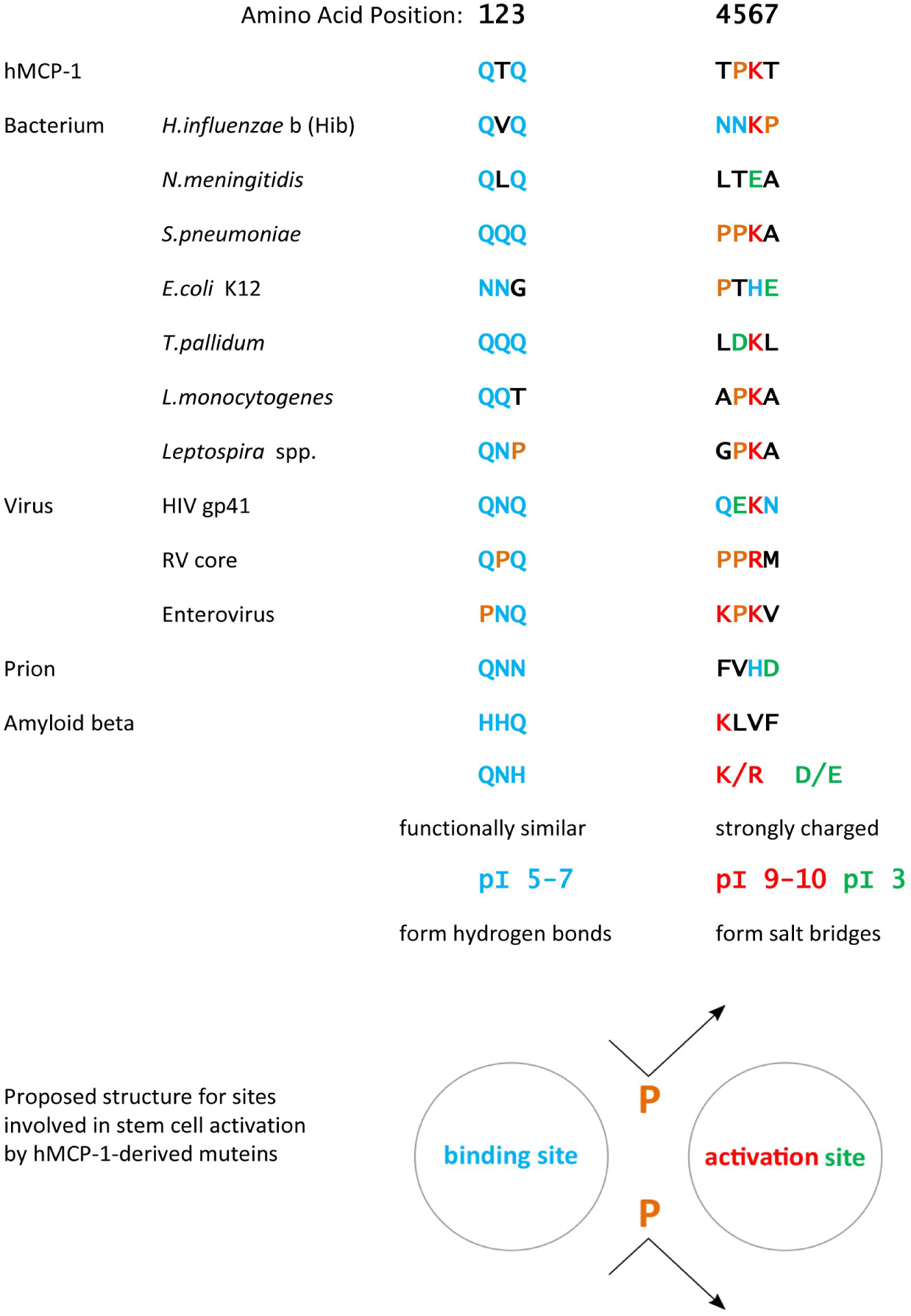
A summary of some of the many homologous muteins of MCP-1 found in bacteria, viruses and spirochetes and in Abeta and prions.

Homologies among the MCP-1 muteins range from 29% (2/7 matches, ex. RV proteins) to 57% (4/7 ex. *L.monocytogenes* p60). Over such short stretches of sequence, these septas are of arguable significance in terms of true homology. Computer generated align plots show no statistically significant relatedness. However, if one allows for charge-conservative substitutions, some of the septas reach 71% similarity with the MCP septapeptide. Despite the relatively low level of sequence conservation, antibody cross-reactivity among the infectious agents expressing septas suggest the possibility of a conserved structure of the septa motif not necessarily reflected in the linear sequence. It was, in fact, the cross-reactivities with a monoclonal antibody raised against RV and other bacterial organisms that cross the BBB to enter the brain that initially suggested an unexpected relatedness.

Taken together the first three residues have pI’s between 5-7 and are functionally similar in their ability to form hydrogen bonds while the majority of the last four residues have pI’s of 3 or between 9-10 and and form salt bridges.

Figure 1 also shows a proposed structure for sites involved in stem cell activation, involving a binding site involving the first three amino acids and an activation site for the last four amino acids. QNH, K/R and D/E are shown as representative amino acids at positions 1-3 and 4-7 respectively. The eponymous prolines would provide a bend in otherwise linear peptides and consequently, differences in the relative positions of these two sites.

We next sought to assess whether the meningitis-associated septas were relevant to disease, by attempting to place them in the the known external portions of the outer membrane protein (OMP) which threads in and out of the surface membrane of Gram negative bacteria.

An external placement of septas ensures maximal chance for contact with, and infection of, the appropriate subset of monocytes destined for entry into the CNS through tight junctions [43,44]. Four of the septa sequences in Figure 1 are shown in red, positioned at or near the tips of the surface exposed loops [45-48].

The relevance of septas to disease has been confirmed in *E.coli* K1. The OMP is a highly conserved, structural molecule among all 17 Gram negative bacteria [49]. When the septa in loop 1 was altered from NNNGPTH to NNAAATH or to NNNGPAA, the bacteria with the altered septas showed a greatly reduced capacity to cause meningitis in a newborn mouse model, with all mice alive at day 7 post inoculation [46].

**Figure 2.**
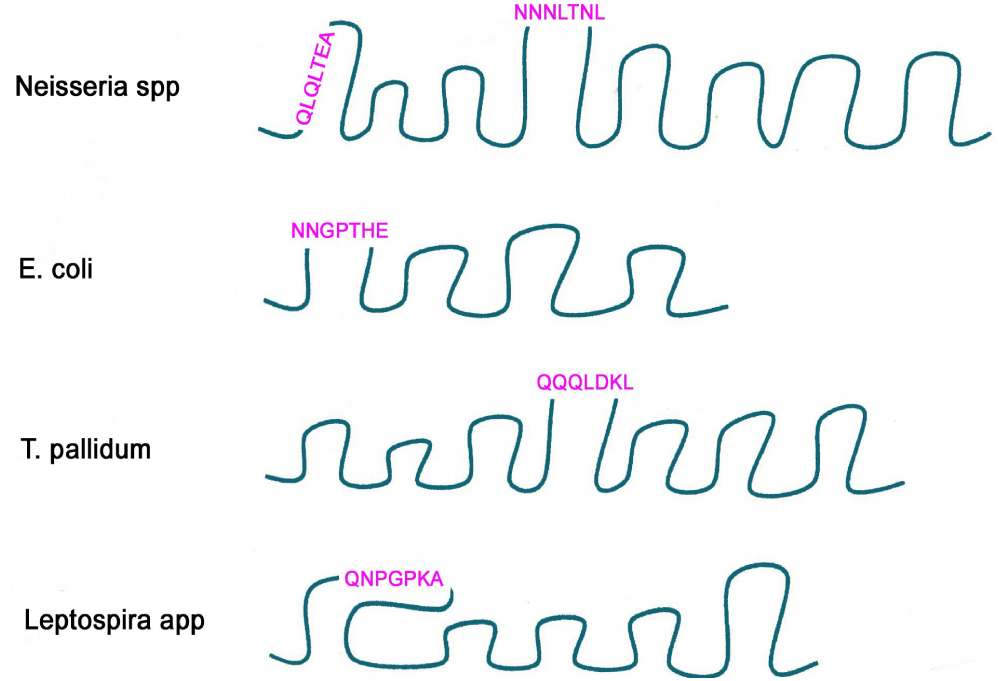
Pathogenic gram negative OMP surface exposed loops.

### Measurement of MCP-1 activity in septas using Ca^+2^ mobilization

We then asked if the septa MCP muteins have chemokine activity by comparing their relative effects on Ca^+2^ mobilization. Some bacterial septas were tested for Ca^+2^ mobilization through receptor-associated heterotrimeric G-protein binding on THP-1 cells.

**Figure 3.**
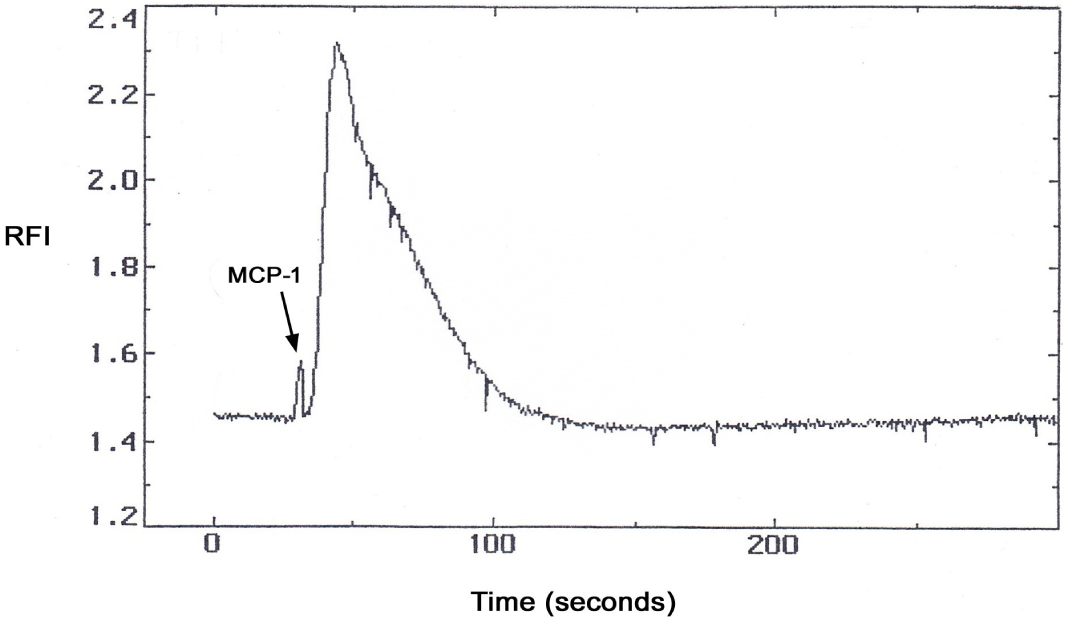
A positive Ca^+2^ mobilization response to hMCP-1 in THP-1 cells. This is the positive control, illustrating the nature of the positive increase in the relative fluorescence intensity (RFI) over time, in response to exposure to MCP-1. The response increase of 0.85 was completed within 100 seconds.

**Figure 4.**
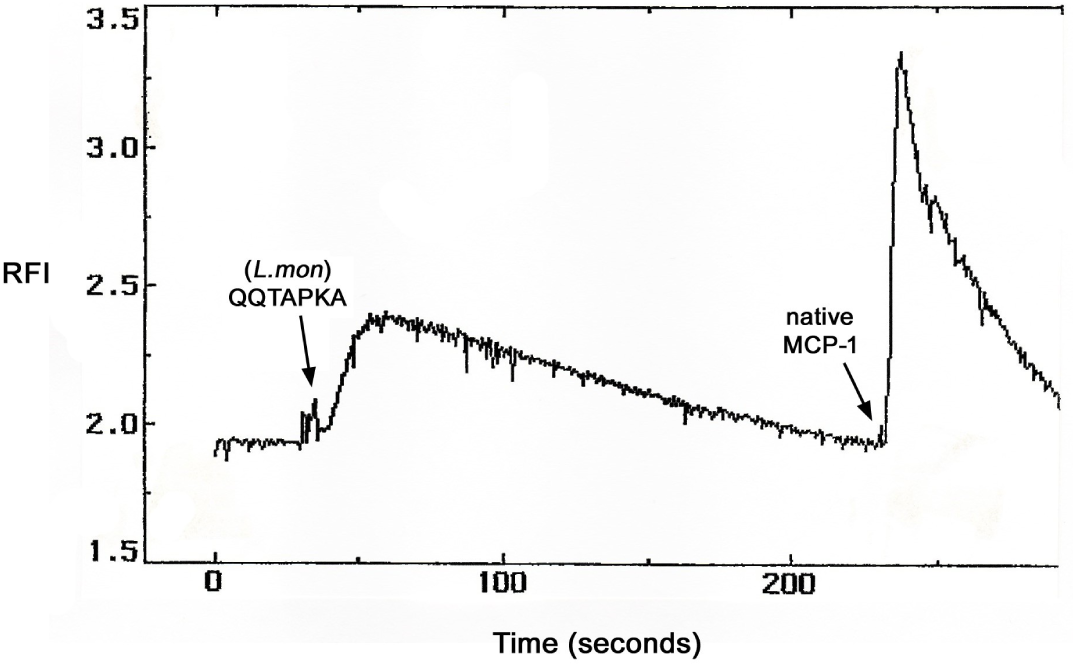
A positive Ca^+2^ mobilization response to the *L.monocytogenes*-derived septa QQTAPKA in THP-1 cells. QQTAPKA elicited an RFI of 0.50, lasting 200 seconds. However, this septa uses a different receptor than the native MCP-1, shown by the lack of subsequent desensitization to MCP-1.

**Figure 5.**
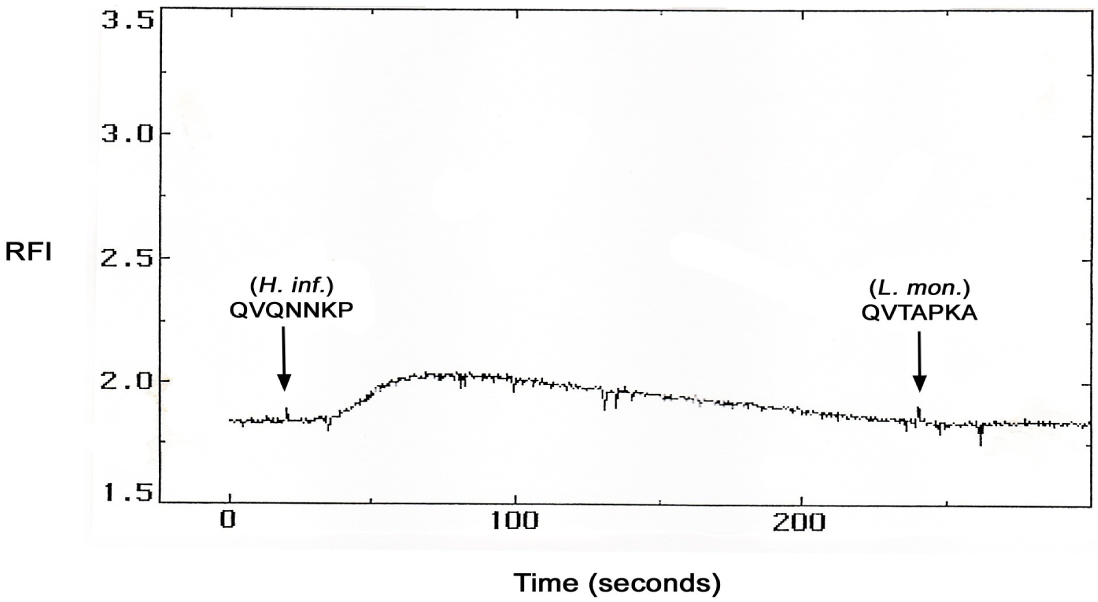
A positive Ca^+2^ mobilization response to the *H.influenzae* type b-derived septa QVQNNKP in THP-1 cells. QVQNNKP elicited a low positive RFI of 0.20, lasting 200 seconds, but blocked activation by the *Listeria*-derived QQTAPKA, indicating that these two septas share a common receptor which differs from that for MCP-1.

**Figure 6.**
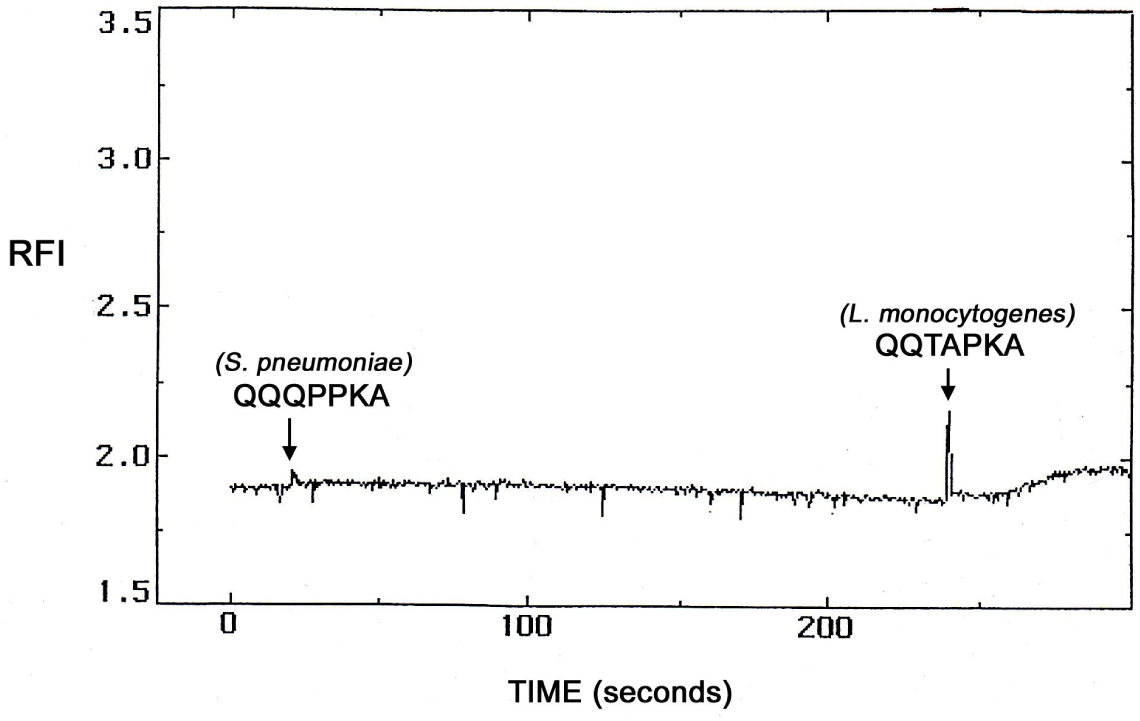
Negative Ca^+2^ mobilization response to the Streptococcal-derived septa QQQPPKA in THP-1 cells. The Streptococcal septa elicited no response itself, but still blocked activation by the *Listeria*-derived QQTAPKA, indicating that these two septa must share at least some portion of a common receptor.

### These findings are summarized in Table 1

**Table 1.** Signalling activity of some bacterial septas in peripheral monocytes

| Organism | septa | $\text{Ca}^{+2}$ flux | Desensitization |
| --- | --- | --- | --- |
| *L.monocytogenes* | QQTAPKA | positive | negative for MCP-1 |
| *H.influenzae* type b | QVQNNKP | positive | positive for QQTAPKA |
| *S.pneumoniae* | QQQPPKA | negative | positive for QQTAPKA |

All randomized septa sequences were negative

Taken together, these data confirm that certain of the free septas acted on their own as mild agonists of Ca^+2^ mobilization. Their signal transduction activity may be mediated through a single (or a single class) of receptor, but the data do not link this with the MCP-1 receptor on THP-1 cells. Further, these data support (1) the hypothesis that the septas represent muteins of the MCP-1 active site for monocyte stem cell activation to macrophages and (2) the notion that infectious organisms may conserve and employ these sequences in order to facilitate their transport through the BBB to infect the CNS.

### RG progenitor cells as indicators of MCP-1 activity

Chemokine activity can be detected to two ways: quantitatively through Ca^+2^ mobilization as previously described, or, by the expression of mitogen-like effects on stem cells in culture. Addition of the intact chemokine or the active site peptide can cause transient differentiation-like events terminating after 72 hours in culture. Activity is scored upon visual examination of the cells at that time.

**Figure 7.**
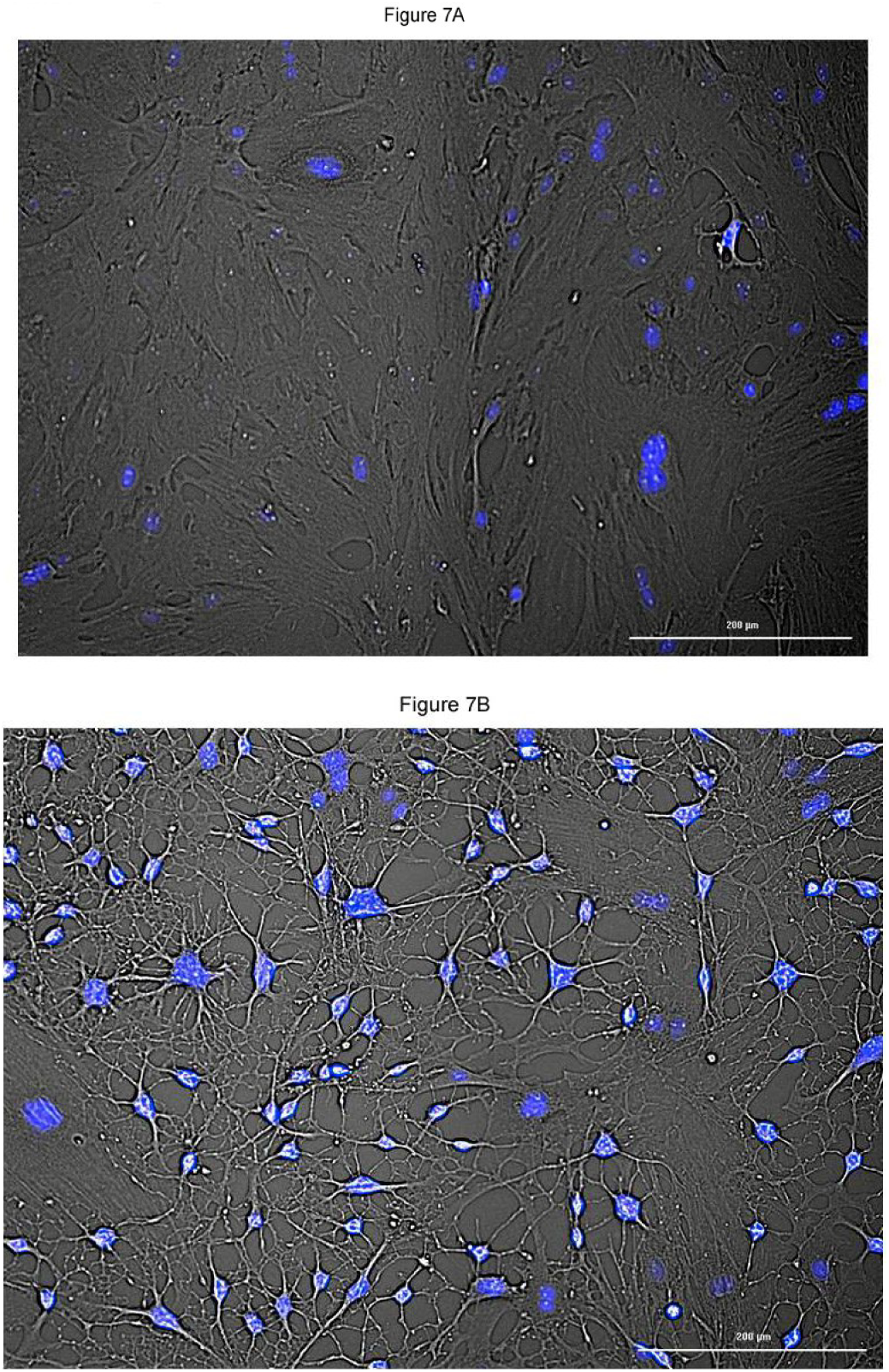
The effect of diBcAMP on monolayer cultures of RG progenitor cells. (A) untreated monolayer (B) RG monolayer treated with 100 uM diBcAMP. Greater than 99% of the cells were transformed into mature glia, with the transient effect greatest at 24 hours. Unstained cells are shown using phase contrast.

Progenitor cells in the third passage after removal from the kainate-treated striata appeared to mature into a diverse population of mature cells.

**Figure 8.**
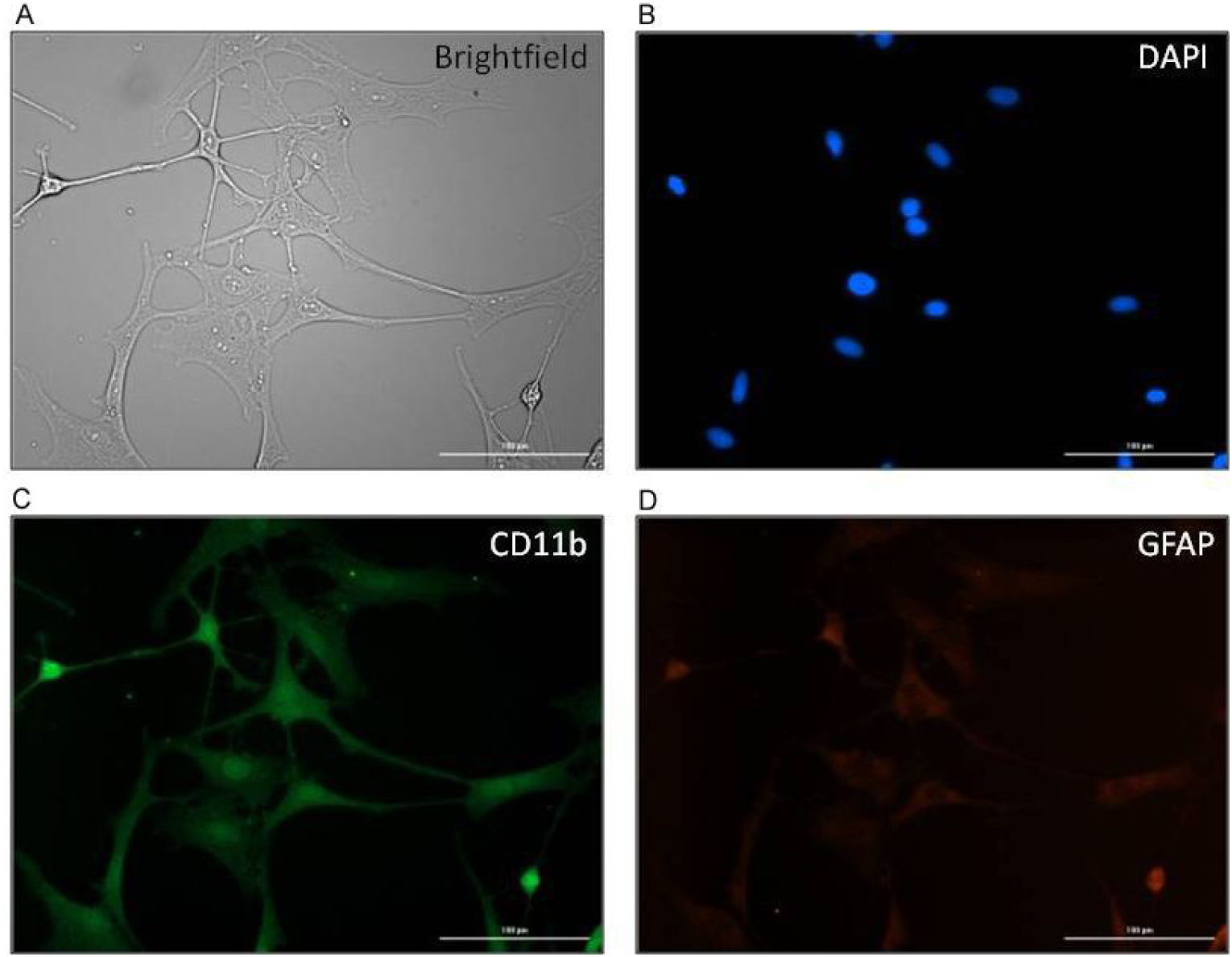
Identification of cells in RG monolayers treated with diBcAMP for 24 hours. One field shown with (A) phase contrast (B) nuclei highlighted with DAPI (C) microglial-specific anti-CD11b antibody staining and (D) astrocyte-specific anti-GFAP antibody staining.

The original characterization of RG cells identified 80% of the population as astrocytes after 35 passages and approximately 15% of the population remained unidentified [39,41]. There was no marker for the identification of microglia at that time. This study, using RG progenitor cells as an early passage, number 3, employed anti-CD11b monoclonal antibody to identify microglia as the predominant cell type following diBcAMP treatment.

### The use of RG cells as indicators of MCP-1 activity using Abeta-derived peptides

Peptides derived from Abeta and synthesized by Kinexus Technologies Inc. (Vancouver, BC, Canada) were tested for their potential MCP-1 effects on RG cell monolayers according to the method of Yankner, et al., 1990 [51]. Since septas may be too short or unstable to form avid bonds with cell surface receptor, the 7’mer was bracketed within slightly longer constructs, in order to maximize stability during the incubation procedure. The septa common to both peptides shown in Table 2 was ^13^HHQKLVF^19^. A randomized sequence was used as a negative control. The peptides tested for maturation activity are shown in Table 2

**Table 2.** Abeta-derived peptides tested for maturation activity using RG progenitor cells in culture.

| Peptide # | Sequence |
| --- | --- |
| 1 | <sup>1</sup>DAEFRHDSGYEV<u>**HHQKLVF**</u><sup>19</sup> |
| 2 | <sup>13</sup><u>**HHQKLVFFAE**</u><sup>22</sup> |
| 3 | FLVKAHEQFH |

where the common septa is shown in underlined, bold, capital letters and where peptide #3 is a randomized negative control for peptide #2

These 3 peptides were dissolved in dimethyl sulfoxide (DMSO) to maintain stability. They were subsequently diluted in PBS. The DMSO solvent caused some distortion of cell morphology at low dilutions up to 10^−3^M,. Therefore, to avoid any interference with the results, RG monolayers were incubated in the presence of peptide concentrations ranging from10^−6^M to 10^−7^M, where there was no detectable DMSO interference, and where chemokine activity is known to be optimally expressed. The results are contained in Table 3, where activity is shown as % conversion, and conversion was scored as an average of 3 readings.

**Table 3.**
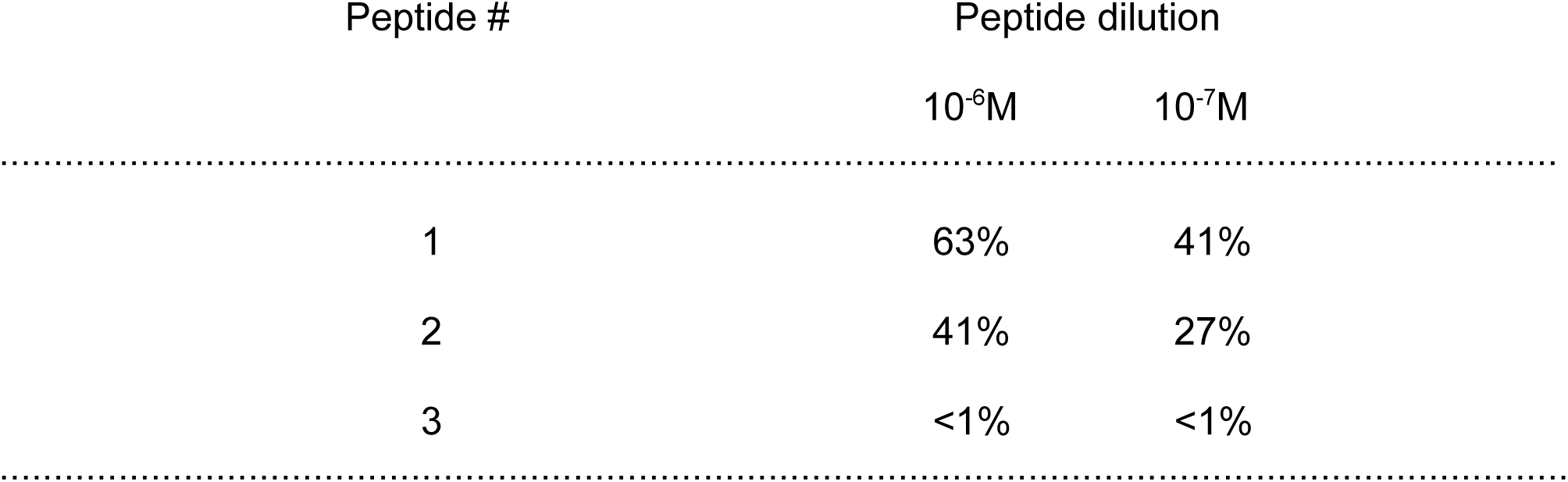
Maturation activity of A-beta-derived peptides on RG progenitor cells in culture

| Peptide # | Peptide dilution |  |
| --- | --- | --- |
| | $10^{-6}$M | $10^{-7}$M |
| 1 | 63% | 41% |
| 2 | 41% | 27% |
| 3 | <1% | <1% |

**Figure 9.**
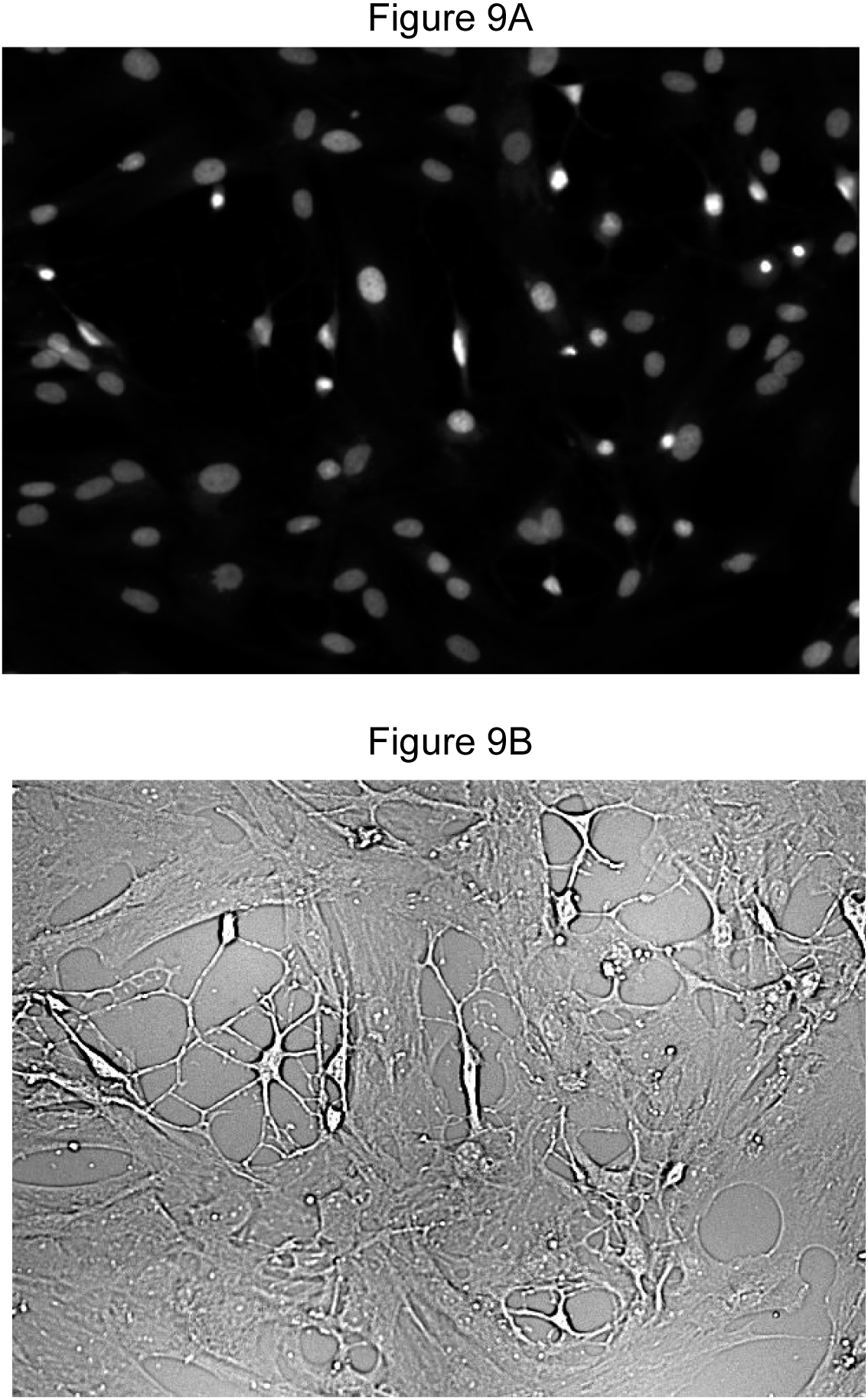
This shows an RG progenitor cell monolayer treated for 72 hours with Abeta peptide number 2 (HHQKLVFFAE) at 10^−7^M. This transient effect was maximal at 72 hours. A. Nuclei counter stained with DAPI B. unstained cells, under phase contrast, showing 27% of the cells in the monolayer exhibiting transformation.

## Discussion

This communication has employed earlier work on RV and meningitis to promote a fresh approach in research into neuodegenerative diseases, such as AD. A new model shown in Figure 10 states that the central event in pathogenesis is the initiation of a chronic, viral or prion infection in the CNS. The resulting membrane damage alerts the brain and triggers the importation and deposition of Abeta, which then expresses hMCP-1 maturation activity on the neighboring glial progenitor cells, leading to an expansion of the mature microglial cell population. Abeta is also known to activate inflammatory signals in microglial cells through Toll-like receptors, adding to the Cell Mediated Immune (CMI) Response to infection. While these glia initiate phagocytic activity, clearing Abeta deposits, infectious organisms and cell debris, after prolonged exposure, they also release a variety of cytotoxic mediators such as TNF-alpha, IL1-beta, chemokines, complement factors, and other neurotoxic factors which can be lethal to surrounding cells [51]. Unfortunately, the overabundance of active microglia generated to combat the original infection would also lead, over time, to the loss of neurons and their synapses culminating in the ensuing dementia.

**Figure 10.**
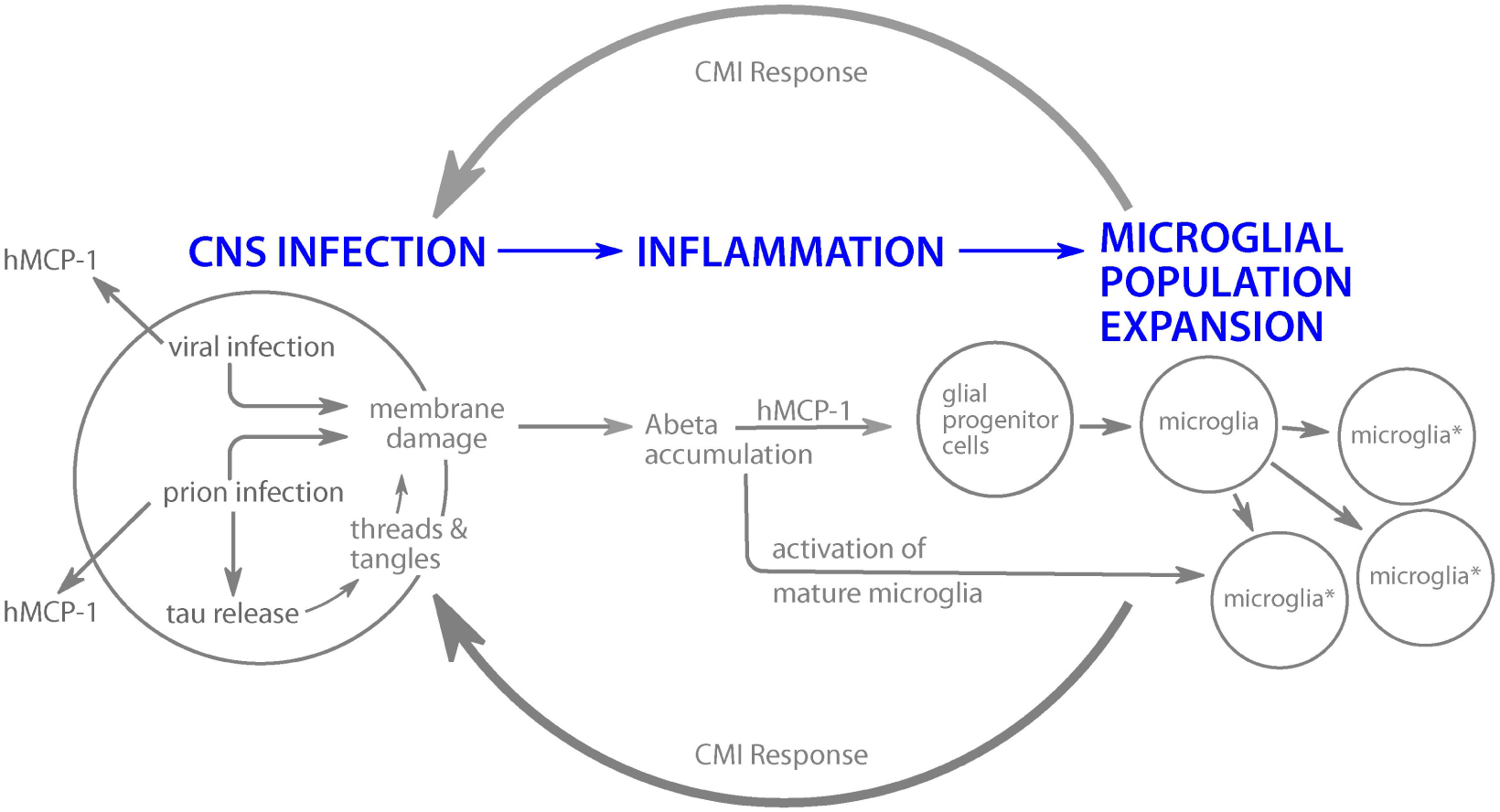
Schematic representation of a new paradigm for neurodegenerative pathology. A CMI response to infection initiated by hMCP-1 activity resident in Abeta is shown. Chemokine activity is also expressed in viral and prion infectious agents, adding to the microglial load. Activated microglia designated as microglia*.

This model provides 3 new concepts to aid in our understanding of neurodegenerative diseases. First, it provides a reasonable explanation for the accumulation of Abeta, as a response to the membrane damage resulting from an active infection. Second, the chemokine activity expressed by Abeta links the amyloid protein directly to the expansion of the microglial population. And, third, it provides new targets for the potential development of therapeutics to treat neurodegenerative diseases.

It must be emphasized that membrane damage occurs only with active viral replication. Chronic viral infection in the CNS may present in two distinct stages, best illustrated by a study of RV infection of RG cells [52-54]. First, infection of progenitor cells, lacking a complete cytoskeleton, led to restricted viral replication. Viral proteins were observed around the nuclear membrane, but no progeny virus particles were detected. Mature virus appeared only after the second stage, when infected cells were treated with diBcAMP and the complete cytoskeleton required for virus assembly and transport to the outer cell membrane was formed. Membrane damage occurs only after active viral replication when mature virus particles reach the outer cell membrane.

The prion protein itself is also thought to be a potential mediator of a variety of neurodegenerative diseases, including AD [29-36]. Prion-infected cells demonstrate a break-down of the cytoskeleton, and the release of the structural protein tau. The further aggregation and hyperphosphorylation of tau ultimately results in the accumulation of abnormal forms of tau aggregates comprising neurofibrillary tangles and threads characteristic of AD pathology. The model presented here also includes a pathway describing tau-related neurodegeneration in two ways. (1) The model predicts that this elevated level of cell membrane damage would alert the brain to import and deposit more Abeta, expressing MCP-1 activity. (2) Just as the Abeta MCP-1 activity is attributed to the septa HHQKLVF,, an homologous MCP-1 septa mutein, QNNFVHD, is found in the prion protein. The model predicts that the prion septa would also express its MCP-1 activity and add to the increasing microglial load.

In light of the parameters of the model presented here, the obvious future strategy for treatment of dementias, such as AD, would be to block the Abeta chemokine activity while leaving the amyloid in place. Plaque would continue to accumulate, but without expression of the MCP-1 activity and concomitant increase in the mature microglial population. It may be insufficient to block the MCP-1 activity of the Abeta alone. The possible contribution to the microglial load by the additional septas expressed by the viruses and/or prions in the chronic infection would have to be considered as well.

This new model refers to chronic viral infection in the widest possible terms, where any virus or prion capable of initiating a chronic infection in the CNS may potentially result in the deposition of Abeta. This list would include many enteroviruses [55], RV [56], measles virus [57], many of the “slow viruses” associated with subacute sclerosing panencephalitis (SSPE) [30], Human Immunodeficiency Virus (HIV) [58], and Herpes Virus, to name just a few. A recent communication has linked human herpesvirus 6A (HHV-6A) and human herpesvirus 7A (HHV7A) with AD across four brain regions from human post-mortem tissue [59]. Hopefully, these data will encourage others to investigate the long term consequences of chronic viral infection in the CNS.

## Summary

Nanomolar concentrations of the Abeta septa ^13^HHQKLVF^19^ were found to express hMCP-1 chemokine activity using rat glial progenitor cells in culture. The data presented here support the hypothesis that the Abeta-derived septapeptide may serve to activate stem cells to increase the number of microglia available to combat chronic infection in the CNS. A new paradigm for neurodegenerative pathology is presented.

Larry Sharma is gratefully acknowledged for his unique and essential contributions, as well as Daniel Bong, Harriett Merks, and Morag Wilson for their excellent technical assistance. It is their steadfast enthusiasm that made this work possible.

